# 3D printed pacifier-shaped mouthpiece for fMRI-compatible gustometers

**DOI:** 10.1101/2021.05.02.442330

**Authors:** David Munoz Tord, Géraldine Coppin, Eva R. Pool, Christophe Mermoud, Zoltan Pataky, David Sander, Sylvain Delplanque

**Affiliations:** Swiss Center for Affective Sciences, University of Geneva, Switzerland; Department of Psychology, University of Geneva, Switzerland; Department of Psychology, Swiss Distance University Institute, Switzerland; Department of Medicine, University of Geneva, Switzerland

**Author notes:** Correspondence should be addressed to: Swiss Center for Affective Sciences, Campus Biotech, Chemin des Mines 9, 1202 Geneva, Switzerland. These authors contributed equally to this work (David Munoz Tord, Géraldine Coppin).

**Keywords:** Mouthpiece, gustometer, fMRI, taste, flavor

## Abstract

Gustometers have allowed the delivery of liquids in fMRI settings for decades and mouthpieces are a critical part of those taste delivery systems. Here we propose an innovative 3D printed mouthpiece inspired by children’s pacifiers that allow participants to swallow while lying down in an MRI scanner. Our results validate the effectiveness of our method by showing significant clusters of activation in the insular and piriform cortex which are regions that have been consistently identified to compute taste processing. We used a large sample (n=85) to validate our method. Our mouthpiece fulfills several criteria guarantying a gustatory stimulus of quality, making the delivery more precise and reliable. Moreover, this new pacifier-shaped design is: simple and cheap to manufacture, hygienic, comfortable to keep in mouth, and flexible to diverse use cases.We hope that this new method will promote and facilitate the study of taste and flavor perception in the context of reward processing in affective neuroscience and thus help provide an integrative approach to the study of the emotional nature of rewards.

## 1 Introduction

Studying the neuronal pathways of chemical senses (i.e., olfaction and gustation) requires special equipment. However, it is relatively easy to make olfactometers (e.g. Coppin, 2020), and the same statement may be even more true for gustometers (e.g. Canna et al., 2019).

The gustometer is a tool specifically designed to deliver liquids. Some gustometers have been used for almost 20 years (e.g. O’Doherty et al., 2002; Small et al., 2003). However, mouthpieces, which are a critical part of the gustatory delivery system (Andersen et al., 2019; Canna et al., 2019), have not been much updated, whereas the number of publications on the topic have kept increasing over the years (see Fig. 1).

**Figure 1:**
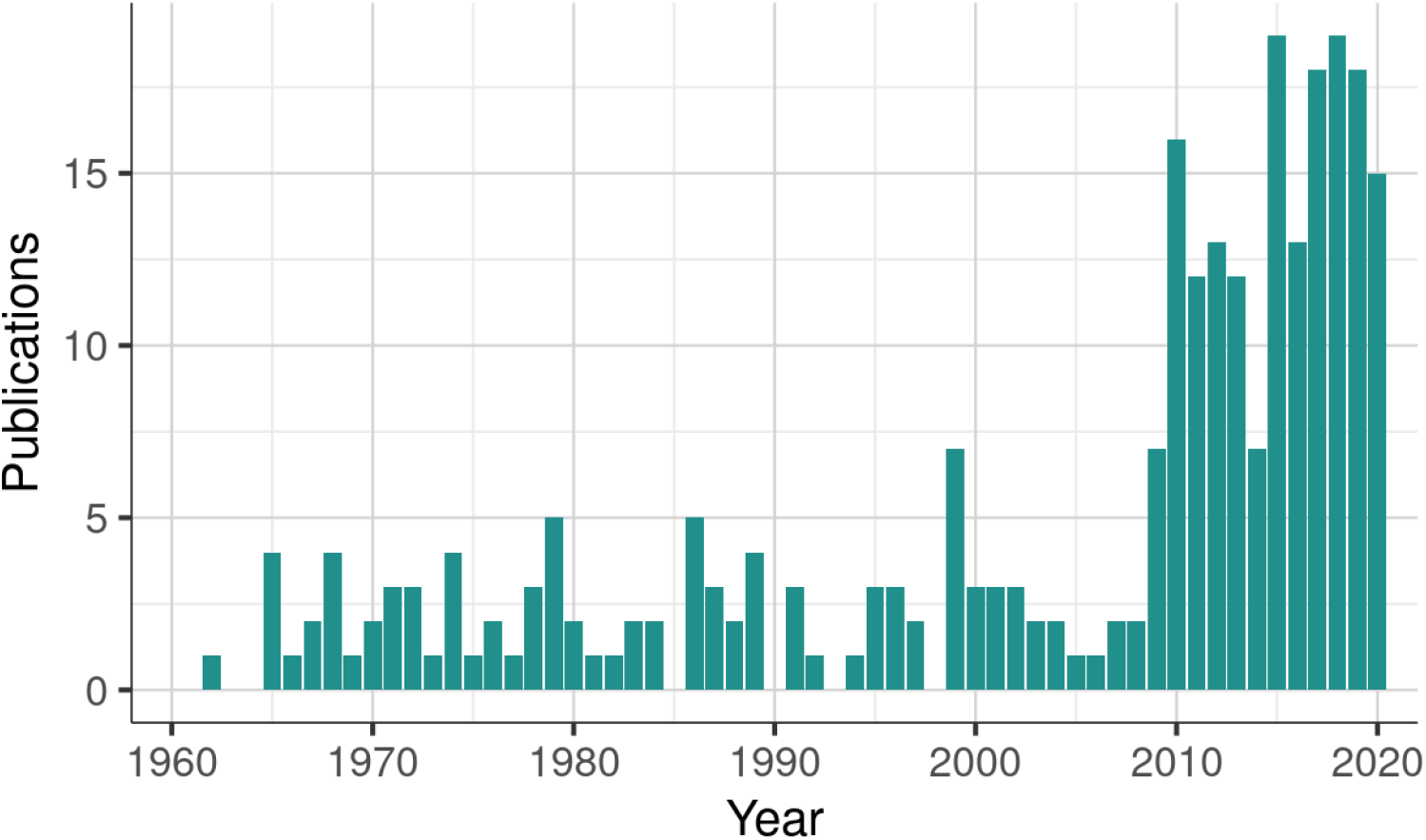
Publications listed on Google Scholar. Results returned when queried with the search terms involving a ‘gustometer’. Results show a clear increase of the number of publication over the years, culminating with over 300 publications on 2020.

Here, we propose an innovative 3D printed MRI-compatible mouthpiece which fulfills several criteria guarantying a gustatory stimulus of quality. First, this new mouthpiece (see Fig. 2) allow participants to swallow while lying down in a scanner, with their heads immobilized in a given position and can remain comfortably in the mouth for an considerable amount of time without requiring any particular effort. Indeed this design –inspired by children’s pacifiers– removes the need for the participant to apply pressure on a ‘biting stick ‘with their teeth and to have to take into account individual dental impressions (e.g. Goto et al., 2015). Second, the mouthpiece permits to deliver up to 8 different liquids in a precise and consistent manner in the center of the tongue. This make it possible to control location delivery, thus minimizing somatosensory variations.

**Figure 2:**
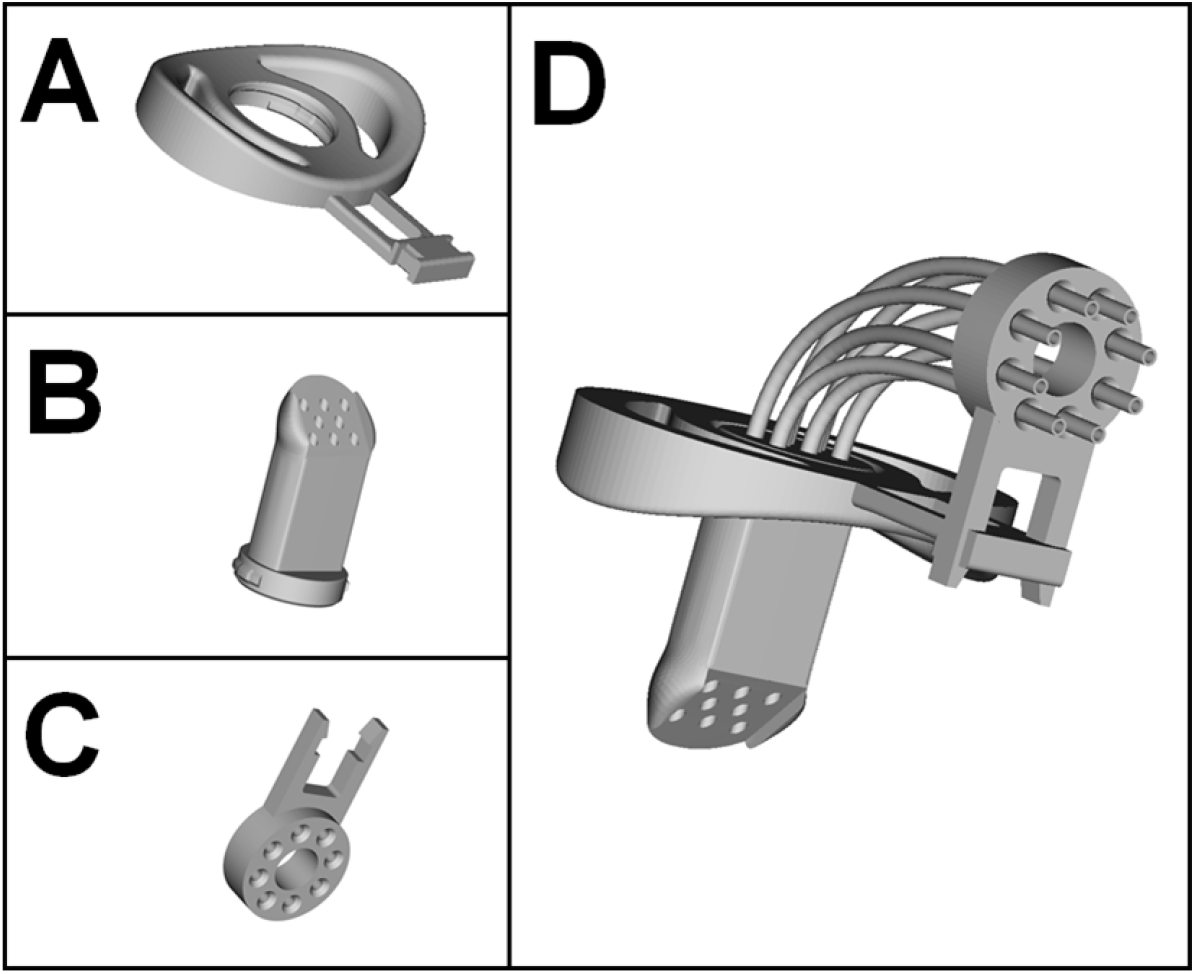
3D representation of the fMRI compatible mouthpiece. Detailed 3D representation of **(A)** the mouth shield, **(B)** the mouthpiece, **(C)** the tube guide and **(D)** the complete mouthpiece assembled with 8 tubes.

**Figure 3:**
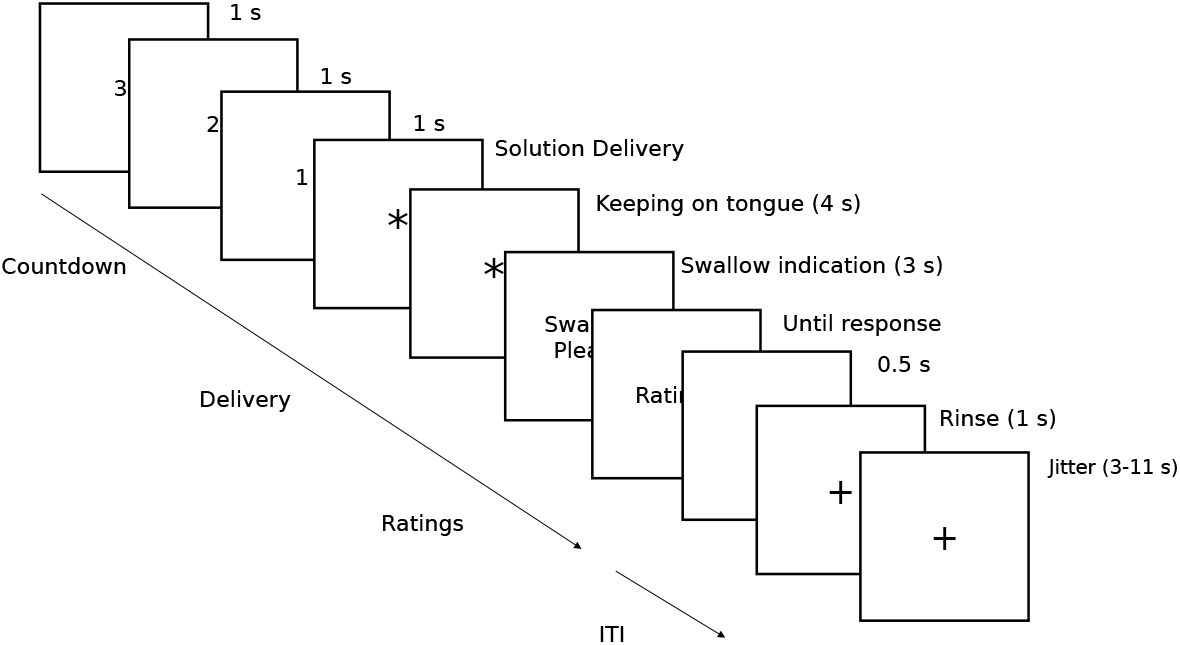
Task procedure. The sequence of the taste reactivity test administered while participants were lying in the scanner. After a brief countdown, the participants were showed an fixation cross followed by the delivery of either a milkshake or a tasteless solution. They were asked to keep the solution on their tongue for 4s and then indicated to swallow it. At this moment they were asked their perceived pleasantness and intensity of the solution. The experimental trials were intertwined with rinse trials to cleanse their palate.

The mouthpiece can be available to any laboratory having access to a 3D printer or could otherwise get them from any 3D printing service company since our plans are made freely available. It can be manufactured in quantity for a very low price (0.5 USD$ of material per piece). This makes it intrinsically hygienic since each participant uses a mouthpiece specially printed for them. Moreover, the printing material can easily be adapted to match different countries’ sanitary regulations. Ours were made out of natural polylactic acid (PLA) compatible with use in contact with food (Conn et al., 1995). Finally, it does not require to modify any pre-existing apparatus and will fit to most gustometer setups seamlessly.

### 1.1 Mouthpiece description

The mouthpiece inspired by children’s pacifier consists of three parts: a mouth shield, an elongated teat and a tube guide. These three pieces are printed separately in natural PLA, a biodegradable plastic made from corn. Other plastics can be used but it remains the responsibility of the researcher to comply with the health standards of the country in which he or she is conducting the experiments with this mouthpiece.

An oval mouth shield (Fig. 2A) holds the mouthpiece comfortably on the lips thanks to its curvature adapted to the morphology of the face. A cylindrical teat (40 mm long x 22 mm diameter) is inserted and clipped on the centre of the mouth shield. This teat receives the tubes at one extremity and directs the liquids to the tongue (Fig. 2B). The part that goes into the mouth and is intended to come into contact with the tongue is bevelled on one side and rounded on the other. This allows for easy contact of the tongue on the teat to deliver drops of liquid comfortably and accurately. Depending on research needs, up to 8 tubes with an external diameter of 2.5 mm (± 0.3 mm) can be inserted into the teat. The last piece is a tube guide (Fig. 2C) that is clipped onto the mouth shield and allows the tubes to be at a 90° angle so that they run along the body of the participant lying on the MRI bed (Fig. 2D). The 3D printing files (stl) that we supply (https://github.com/munoztd0/Mouthpiecegusto) include seven versions with a diameter of 2.5 mm ± 0.3 mm in steps of 0.1 mm. All these versions make it possible to choose the parts that fit together best according to the 2.5 mm tubes used by the researchers and allows to adjust for different types of liquid or viscosity levels.

## 2 Method

### 2.1 Participants

This study was part of a larger experiment that was related to a different study question (reward processing) in which 97 right-handed participants were recruited. The study was approved by the Swissmedic ethical committee. All participants gave written informed consent and received 200 Swiss francs for their participation to the whole session. In total, 12 participant were discarded from the analysis because of missing or incomplete data (5 MRI and 7 behavioral).

We report data on the 85 remaining participants (55 female; mean age, 37.3 ±12.4; min–max, 18–67 years). No predetermined sample size was estimated via statistical methods. None of the participants have reported having any kind of olfactory disorder. All participants were asked to have fasted overnight to our experiment that happened in the morning.

### 2.2 Preparations

Milkshake preparations were made from a mix of milk (300 g) and ice cream (60 g) for a total of 71 kcal/100 g. Potassium chloride (KCl, 1.8 g) and sodium bicarbonate (NaHCO3, 0.21 g) were diluted in 1L of distilled water to recreate an artificial tasteless saliva solution. This main solution was then used to create less concentrated preparations to be able to further match each individual’s perception of a tasteless solution. In total, there were 4 different tasteless concentrations (1/1, 3/4, 1/2 and 1/4) and 3 flavors of milkshake (strawberry, chocolate or vanilla). Each participant was asked which flavor of milkshake they preferred as well as which saliva solution they found more neutral. This two solution where then used as the main two stimuli for the rest of the experiment.

### 2.3 Gustometer

Single channel syringe pumps (Chemyx OEM) were used to achieve high flow control. Two syringes of up to 60 mL were connected via 8 meters length polyurethane food grade tubing (external diameter = 4 mm, inner diameter = 2.5 mm) to 1 meter length food grade PTFE tubing (external diameter = 2.5 mm, inner diameter = 1.9 mm) and then to the mouthpiece at a delivery rate of 1 mL/s. The syringe pumps were connected to a 16-port RS-232 rackmount device server (Moxa, Nport 5610) and then controlled via TCP/IP using specific C libraries designed for stimulus presentation software (E-prime, Matlab or python). While it is out of the scope of this paper but readers can refer to (Andersen et al., 2019; Canna et al., 2019; Haase et al., 2007) for detailed instructions on how to setup a MRI-compatible gustometer.

### 2.4 Taste Reactivity Task

An taste reactivity test was administered while participants were lying in the scanner. The task consisted in the evaluation of the perceived pleasantness and intensity of two different stimuli: a milkshake and a tasteless solution. We chose individually adjusted tasteless solution as control stimulus instead of plain water because water has been shown to have an inherent taste (Bartoshuk et al., 1964). Each trial consisted on the administration of 1 mL of the solution and the delivery order of the two conditions were randomized within each participant. Participants were asked to keep the solution on their tongue for 4s before swallowing to avoid adding movement noise to the fMRI response. The experimental trials were intertwined with rinse trials to cleanse the participants’ palate with 1 mL of water.

All 40 evaluations (20 per solution) were done on visual analog scales displayed on a computer screen. Participants had to answer through a button-box placed in their hand. The visual scales for the intensity report ranged from “not perceived” to “extremely intense”; and from “extremely unpleasant” to “extremely pleasant” for liking ratings.

### 2.5 Data Acquisition

The collection of the responses were controlled by a computer running MATLAB (version R2015a; MathWorks, Natick, USA). The presentation of the stimuli was implemented using Psychtoolbox (version 3.0). The acquisition of the neuroimaging data was performed via a 3 Tesla Magnetom TrioTrim scanner (Siemens Medical Solutions, Erlangen, Germany) supplied with a 32-channel head coil following a gradient echo (GRE) sequence to record Blood-Oxygen-Level-Dependent (BOLD) signal. We recorded forty echo-planar imaging (EPI) slices per scan with a isotropic voxel size of 3 mm. Our scanner parameters were set at: echo time (TE) = 20 ms, repetition time (TR) = 2000 ms, field of view (FOV) = 210*×*210*×*144 mm, matrix size = 70*×*70 voxels, flip angle = 85°, 0.6 mm gap between slices.

Besides structural whole brain T1-weighted (T1_*w*_) images (isotropic voxel size = 1.0 mm), dual gradient *B*_0_ field maps (Fmaps) were also acquired for each participant to deal with distortions caused by inhomogeneity in the static-field.

### 2.6 Preprocessing

We combined the Oxford Centre’s *FMRIB* Software Library (FSL, version 4.1; Jenkinson et al., 2012) with the Advanced Normalization Tools (ANTS, version 2.1; Avants et al., 2011) to create a pipeline optimized for the preprocessing of our neuroimaging data (see Suppl. Fig. S1).

A challenge of fMRI gustometry is that BOLD signal is highly prone to movement artifacts and thus the swallowing of liquid solutions while lying down produces significant deglutition artifacts. To offset this loss of signal we followed (Griffanti et al., 2017) rigorous protocol for fMRI ICA-based artifact removal (e.g. motion, susceptibility or blood flow in arteries). Field maps were then applied to correct geometric distortion and ANTS was used to diffeomorphically co-register the preprocessed functional and structural images to the Montreal Neurological Institute (MNI) space, using nearest-neighbor interpolation and leaving the functional images in their native resolution. Finally, we applied a spatial smoothing of 8 mm full width half maximum.

### 2.7 Data Analysis

Statistical analyses of the behavioral data were performed with R (version 4.0; R Core Team, 2019). We report Cohen’s *d*_*z*_ and their 95%*CI* as estimates of effect sizes for the paired *t* tests (Lakens, 2013) together with a Bayes factor (*BF*_10_) quantifying the likelihood of the data under the alternative hypothesis relative to the null hypothesis Morey et al. (2015).

The Statistical Parametric Mapping software (SPM, version 12; Penny et al., 2011) was used to perform a random-effects univariate analysis on the voxels of the image times series following a two-stage approach to partition model residuals to take into account within- and between-participant variance (Holmes and Friston, 1988; Mumford and Poldrack, 2007).

We specified a subject-level general linear model (GLM) for each participant and added a high-pass filter cutoff of 1/128 Hz to eliminate possible low-frequency confounds (Talmi et al., 2008). Each regressor of interest was derived from the onsets and duration of the stimuli and convoluted using a canonical hemodynamic response function (HRF) into the GLM to obtain weighted parameter estimates (*β*). The subject-level GLM consisted of six regressors: (1) the trial, (2) the reception of the milkshake solution, (3) the reception of the tasteless solution, (4) water rinsing, (5) question about solution pleasantness and, (6) intensity. Group-level statistical *t*-maps were then created by combining subject-level estimated beta weights (Milkshake>Tasteless) and residuals.

We used the AFNI’s *3dFWHMx* function to estimate the intrinsic spatial smoothness of our xyz dimensions that we inputed in the new *3dClustSim* function (Cox et al., 2017) to create–via Monte Carlo simulation–a cluster extent threshold corrected for multiple comparisons over the whole brain at *p* < 0.05 for a height threshold of *p* < 0.001. We report the minimum extent threshold, the peak MNI coordinates, and the number of consecutive significant voxels at *p* < 0.005 within the cluster (*k*). Finally, we display the statistical *t*-maps of our group results of the Milkshake > Tasteless contrast surviving cluster-level correction overlaid on a high-resolution template in MNI space.

## 3 Results

We analyzed the taste intensity ratings during our task using a paired *t* tests with two conditions (milkshake or tasteless). As expected, participants rated the milkshake solution as more intense than the tasteless solution (*F*_(1,84)_ = 153.81, *p* < 0.001, 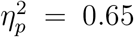, 90% CI = [0.54, 0.72], *BF*_10_ = 2.35 *×* 10^23^, see Fig. 4A).

**Figure 4:**
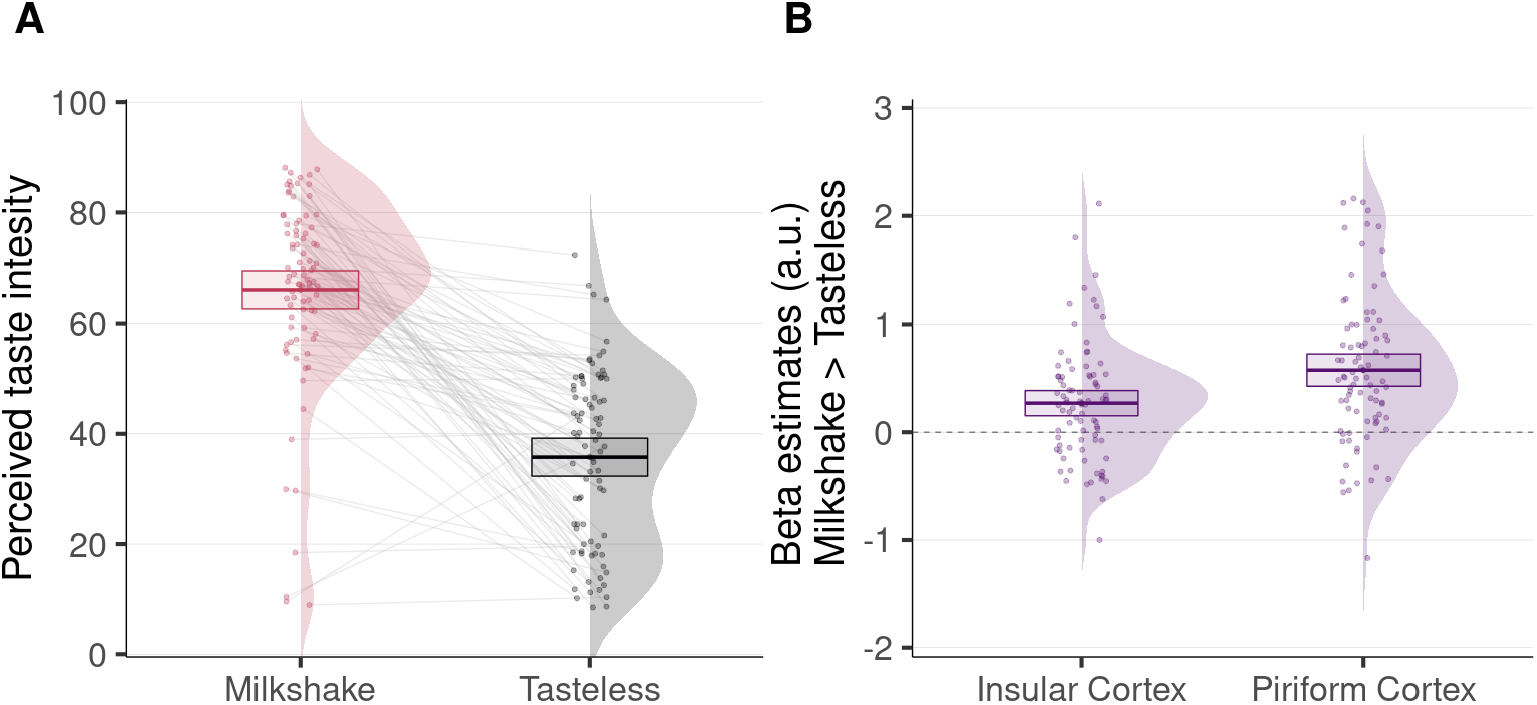
Behavioral and fMRI results. **(A)** Individual estimates, densities and overall mean of perceived taste intensity of the milkshake and the tasteless solutions. **(B)** Individual beta estimates, densities and overall means of the Milkshake > Tasteless contrast across participants during taste delivery extracted from voxel clusters within the insular and piriform cortex. Error bars represent 95% CI (n = 85).

We report the results from our group-level analysis using a height threshold of *p* < 0.005, with a minimum cluster extent threshold corrected for multiple comparisons at *p* < 0.05 of 123 voxels. For the taste reactivity task, the pleasant solution (Milkshake > Tasteless) activated the primary olfactory (piriform) cortex bilaterally (right: MNI [*xyz*] = [−22 −3 −14], *k* = 282; left: MNI [*xyz*] = [21 −6 −14], *k* = 149), the primary gustatory (insular) cortex (left: MNI [*xyz*] = [21 −6 −14], *k* = 149), and the primary somatosensory (parietal operculum/postcentral gyrus, see Fig. 5 and Suppl. Table S1 for more details).

**Figure 5:**
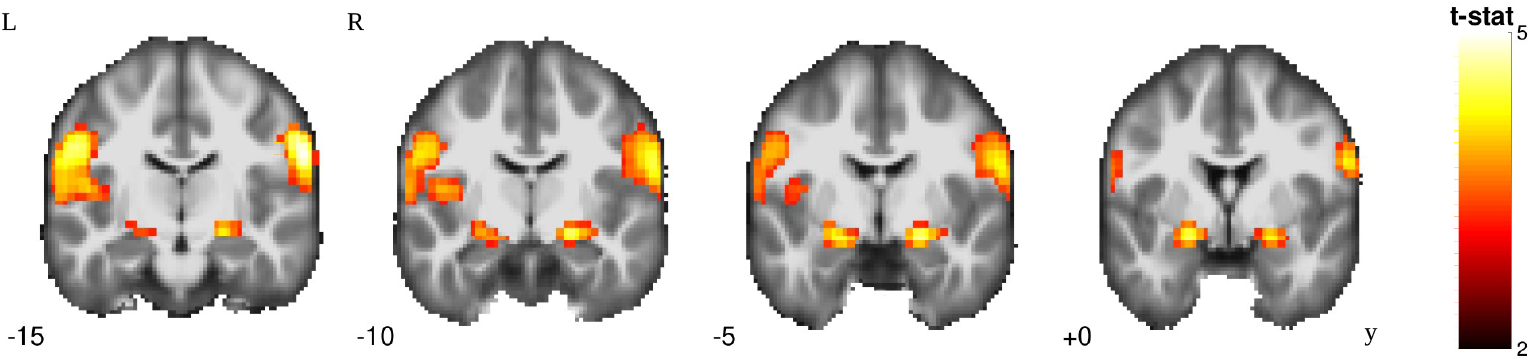
Neural correlates of taste. Regions which the BOLD signal positively correlates with the magnitude of the contrast Milkshake > Tasteless (n = 85). Statistical *t*-maps are shown with a threshold of *p* < 0.001 and a minimum cluster extent threshold (corrected for multiple comparisons) of 123 voxels.

We also computed observed power calculations within two regions, namely the insular and piriform cortex. To achieve that we extracted the averaged betas values from within these two regions and calculated their standardized effect size (*d*_*z*_). We report a *d*_*z*_ = 0.41 for the insula and *d*_*z*_ = 0.56 for the piriform, this allowed us to estimate that to achieve a 90% power at *α* = 0.05 one would need 53 or 29 participants, for the insula or the piriform respectively, to reproduce these results (see Suppl. Fig. S2).

## 4 Discussion

In this paper, we presented a 3D printed MRI compatible mouthpiece for the study of human taste and flavor perception in fMRI settings. After describing this mouthpiece, we reported the results of 3 Tesla fMRI study and, as illustrated by our findings, this mouthpiece allows to obtain an effective measure of brain related activity during the consumption of gustatory stimuli.

We provide results from a large sample that both demonstrate the effectiveness and validity of our procedure by showing significant clusters of activation within the same regions that have been reported throughout different meta-analyses on taste (Yeung et al., 2017) and olfaction (Seubert et al., 2013).

We found clear activations of: (i) the middle insular cortex which was no surprise since this region has consistently been identified as the the human primary gustatory cortex (Buck and Bargmann, 2000; Small and Faurion, 2015), (ii) the parietal operculum/postcentral gyrus which has been reported to be the primary cortex for oral somatosensory representation in humans (Boling et al., 2002) and, (iii) the anterior medial temporal lobes –including the hippocampal formation and the amygdaloid complex– which have also both been revealed to play a crucial role in food intake (Coppin, 2016; Davidson et al., 2009; Petrovich, 2011).

We however encountered some limitations that should be addressed. First, some participants reported that a 40 mm long mouthpiece was a bit too long and thus uncomfortable. This can easily be alleviated by printing a shorter mouthpiece in those cases. We also tried to extend our setup to a non-MRI contexts were participants would be seated in a upright position. It appeared that the liquids would not flow as consistently and precisely that they did in a lying position and would suggest that the prototype would have to be modified for such contexts.

In a few cases and during intensive use, we also noticed that the plastic could become porous, so that the joints between the tubes and the teat were no longer perfectly sealed. As a result, some participants reported that rinsing liquid had run down their cheeks. However, this did not prevent the stimuli from being sent, but it is something that the researchers could monitor. One option might be to choose a less porous plastic that is still within the country’s legislative constraints on plastics permitted for food contact.

Moreover we think it is important to tell the participants to place their tongue in such a way as to let the solutions flow without blocking the teat to deliver drops of liquid comfortably and accurately.

## 5 Conclusion

The main advantages of this mouthpiece are its low cost, flexibility, ease to produce and fMRI-compatible design. Any lab with access to an 3D printer can make one or could otherwise get them from any 3D printing service company since our plans are made freely available. But most importantly, it is flexible and can be modified to any particular case. It can easily match different countries’ sanitary regulations or be adjusted for different types of liquid or viscosity levels. It also does not require to modify any pre-existing apparatus and will integrate to most gustometer setups without any additional work.

We think that this new method will promote the use of primary rewards (i.e. milkshakes) instead of more widely used secondary rewards (e.g. food pictures) to measure hedonicity. This is extremely important because, for one part, it allows comparison with the animal literature on innate food rewards; but also avoids reward type-dependent neural circuits of secondary rewards (Nakamura et al., 2020; Sescousse et al., 2013). Moreover, taste consumption can induce an affective experience by itself rather than a representation of the affective experience (i.e. pictures of food) which is a crucial property for the proper study of processing rewards (Pool et al., 2016, 2021).

Affective neuroscience would benefit from the inclusion of more studies in olfaction and taste using primary rewards. This would provide the means for an integrative approach to study the emotional nature of reward (Nummenmaa and Sander, 2020).

## Data and materials availability

Unthresholded statistical *t*-maps are available at the Neurovault platform (neurovault.org/images/442236/). Computer code used for producing the mouthpiece as well as preprocessing and analyzing the data is available in a publicly hosted software repository (github.com/munoztd0/Mouthpiece_gusto).

## Credits

DMT analyzed the data. ERP help with the validation. CM designed the apparatus. ZP provided the participants. GC and DS designed the experiments and SD managed the project. DMT, GC and SD wrote the first draft. All authors reviewed and approved the final manuscript.

## Acknowledgments

The authors would like to thank Alain Hugon for his major contribution in the early stages of the design of the pacifier-shaped mouthpiece and Asli Erdemli for her useful comments from the data acquisition and Lavinia Wuensch for her work on the data preprocessing. We also thank all the people from the Perception and Bioresponses Department of the Research and Development Division of Firmenich, SA for their precious advice and their theoretical and technical competences.

