## Supplementary Material for "3D printed pacifier-shaped mouthpiece for fMRI-compatible gustometers"

### A. Supplementary Material

#### Artifact removal

Two researchers independently hand classified the IC for one task (~150 IC) from each participant into two categories: 'clear artifact' (e.g. motion, susceptibility or blood flow in arteries) or 'potential signal'. Labels were then compared between the two judges where each discrepancy was discussed until an agreement was reached (inter-rater reliability = 93%). Manually classified components obtained by this process were used to train a classifier using random forest machine learning algorithm Breiman (2001). Leave-one-out testing—where we iteratively left one participant out of the training and tested the classifier accuracy on the left out participant—at the optimal sensitivity (threshold = 5) resulted in a median 94% true positive rate (i.e. the percentage of 'true signal' accurately classified). Consequently, we applied *FIX* to automatize the denoising of our BOLD signal Salimi-Khorshidi et al. (2014).

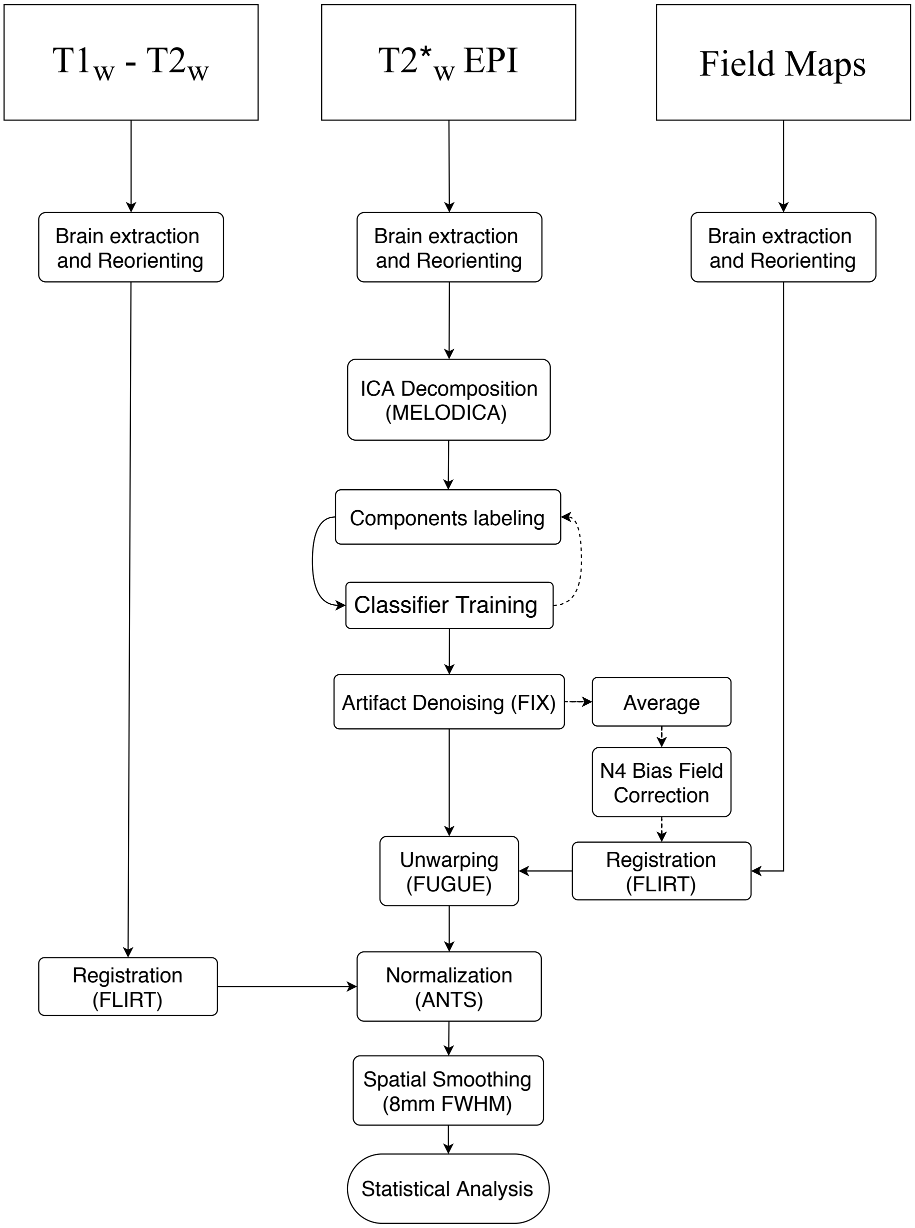


Figure S1: **Flow chart representation of the preprocessing pipeline**. Costum build pipeline to improve the removal of deglutition artifacts in functional MRI.

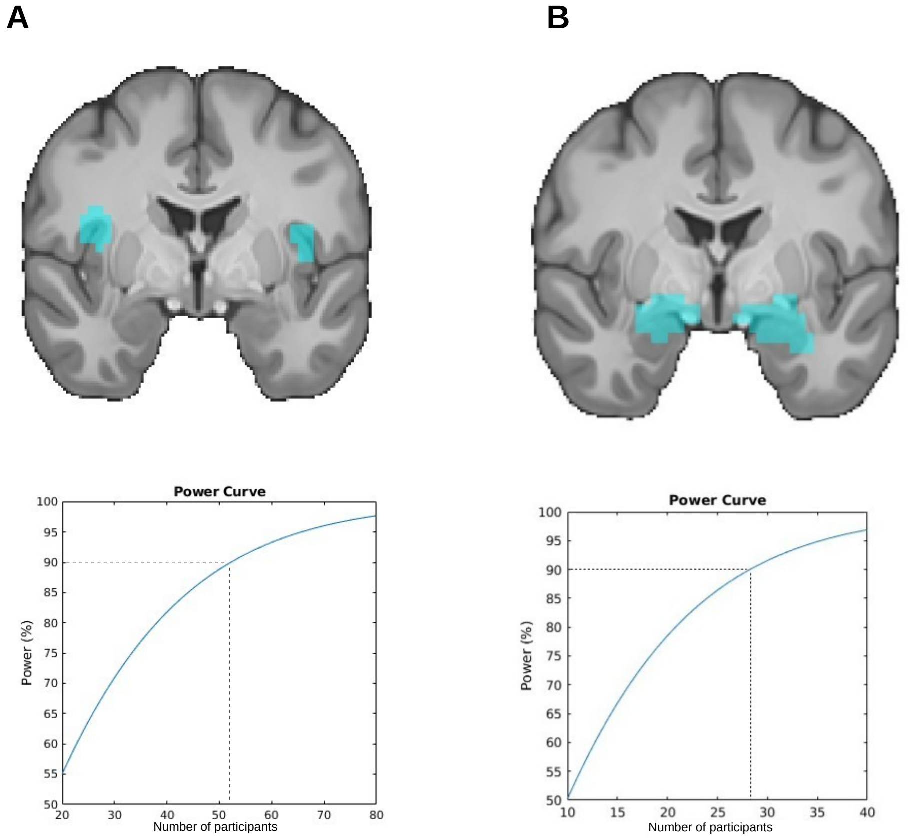


Figure S2: **Observed power**. **(A)** One would need 53 participants to reproduce our results within the insular cortex with a power of 90% and an $\alpha = 0.05$. **(B)** One would need 29 participants to reproduce our results within the piriform cortex.

Table S1: Summary Results of BOLD Activations in the Taste Reactivity Tests

| Region | Laterality | Extent | $t(82)$ | Coordinates | | |
|---|---|---|---|---|---|---|
| | | | | x | y | z |
| Parietal operculum / Postcentral gyrus | L | 532 | 5.19 | -55 | -21 | 36 |
| Insular cortex | L | 532 | 4.15 | -49 | -12 | 11 |
| Parietal operculum / Postcentral gyrus | R | 327 | 4.96 | 63 | -15 | 29 |
| Piriform Cortex / Amygdala | R | 282 | 4.78 | -22 | -3 | -14 |
| Parahippocampal gyrus | L | 282 | 4.77 | -13 | -36 | -7 |
| Piriform cortex / Amygdala | L | 149 | 4.73 | 21 | -6 | -14 |
| Parahippocampal gyrus | R | 149 | 3.72 | 15 | -30 | -7 |

*Note.* Thresholding $t(82) > 3.19$, $p < 0.001$, and minimum cluster level simulation extent for multiple comparison correction at $p < 0.05 = 123$. Table shows all local maxima separated by more than 20 mm. Coordinates are expressed in the Montreal Neurological Institute (MNI) space in the left-right, anterior-posterior, and inferior-superior dimensions, respectively.
